# Cross-species neural co-culture uncovers metabolic signatures of cellular crosstalk

**DOI:** 10.64898/2026.07.31.741820

**Authors:** Katherine Rickelton, Rithvik Sandiri, Jessica Roy, Aliyah Dalier, Courtney C. Babbitt

## Abstract

Primates are distinguished by large brains relative to body size, with humans showing the greatest expansion. This increase in brain size evolved alongside advanced cognitive abilities as well as an elevated energetic demand. Importantly, allometric scaling alone does not explain this increased metabolic requirement, suggesting that other cellular mechanisms may be driving the unique energetic capacity of the human brain. Brain metabolism is critical for neurological function by providing the energy necessary for neuron firing. Much of metabolism in the brain is carried out by astrocytes: a type of glial cell that have long been viewed as passive support cells for neurons. More recent research has highlighted the unique roles of astrocytes in many critical neurological processes; however, it is less understood how astrocytes differ among species. To better characterize this, we developed a cross-species co-culture model of astrocytes and neurons from human or chimpanzee-derived iPSCs. This co-culture system allowed us to assess cell-type specific effects as well as species-specific differences in cellular interactions that may be driving overall differences in brain metabolism. We conducted single-cell RNA-sequencing as well as Seahorse XF Mitochondrial Stress tests and observed that human neural co-cultures are more metabolically active than chimpanzee neural co-cultures. Cross-species co-culture systems also highlight that astrocytes are driving major species differences in metabolism, whereas neurons are highly responsive to astrocytic activity. We conclude that both neurons and astrocytes have evolved differently across primates, and that metabolic interactions between these cell types are key contributors in primate brain evolution.

## Introduction

Despite comprising only ∼2% of total body mass, the human brain consumes ∼20% of the body’s glucose supply - a significant increase relative to the <10% observed in nonhuman primates ^1,2^. While the primate lineage is defined by having large brains relative to body size, allometry alone does not fully explain this increased energetic demand in humans ^3–5^. Rather, factors beyond brain size, including cellular interactions and metabolic specialization, likely drive these changes in humans^6,7^.

One proposed hypothesis attributes this increased energetic demand to the greater number of neurons found within the human brain ^8^. Neurons, the electrically active cells in the brain, require significant metabolic support to facilitate communication between cells^9,10^. Providing that metabolic support are glial cells, a non-neuronal class of cells that are not electrically active but, as the most abundant cells in the brain, have important roles in maintaining brain homeostasis ^10–12^. With an increase in neuron number comes a similar expansion in glial cell content, and the glia-to-neuron ratio has gradually increased throughout the primate lineage - ranging from 0.6 in some new world monkeys to 1.65 in humans ^8,13–15^. This data suggests that glia may be well positioned to influence the heightened metabolic demands of the human brain, however, the extent to which their functional properties differ across species remains poorly understood.

Astrocytes, a type of glial cell, provision metabolites to neurons in order to support energetically expensive synaptic processes ^11,16–18^. Astrocytes are uniquely characterized by high aerobic glycolytic activity (increased glycolysis and limited potential for oxidative phosphorylation) whereas neurons typically favor oxidative phosphorylation for high levels of ATP production ^19–21^. With this, astrocytes are thought to respond to neuronal signals associated with increased synaptic activity by increasing glucose uptake and subsequent aerobic glycolysis^21,22^. These astrocyte–neuron interactions form a metabolic feedback loop in which astrocytes support synaptic activity, and synaptic signaling reciprocally regulates astrocytic function.

Astrocytes also display striking evolutionary divergence: human astrocytes are significantly larger and form more complex branching patterns than observed in rodent models^23–25^. Previously collected transcriptomic data reports that twice as many genes were differentially expressed between humans and chimpanzees in astrocytes compared to neurons, many of which were found to be enriched in metabolic functions ^26^. Single-cell RNA sequencing (scRNA-seq) of post-mortem brain tissue also showed that astrocytes and oligodendrocyte progenitors displayed more transcriptomic differences in the human lineage compared to neurons ^27^.

Nonetheless, cell-type-specific analyses of primate brains remain limited, primarily due to the scarcity and variable quality of post-mortem tissue ^28,29^. Advancements in induced pluripotent stem cell (iPSC) technology help to solve this issue of sample availability and also offer the advantage of greater control over the extracellular environment, maintaining similar growth conditions over time and across species ^29^. Much of the work utilizing non-human primate iPSCs has been conducted using a 2D monoculture system, which fails to recapitulate the interaction networks between multiple cell types ^22^. A co-culture model of multiple cell types (grown in 2D or in 3D) better allows for assessment of how multiple cell types in the brain interact with each other, and further how these interactions may differ among species ^30–33^.

To gain deeper insights into the cellular mechanisms underlying primate brain evolution, we developed a co-culture model of iPSC-derived astrocytes and neurons from humans and chimpanzees (Figure 1). Using this system, we explored both species-specific and cell-type-specific contributions to brain metabolism via a multi-species set up of neurons and astrocytes. We conducted Seahorse XF Mitochondrial Stress Tests to measure real-time metabolic activity in live cells and employed single-cell RNA sequencing to assess transcriptional differences across co-culture conditions. Together, these approaches provide an integrated view of how astrocyte-neuron interactions may have evolved to support the heightened energetic demands of the human brain.

**Figure 1.**
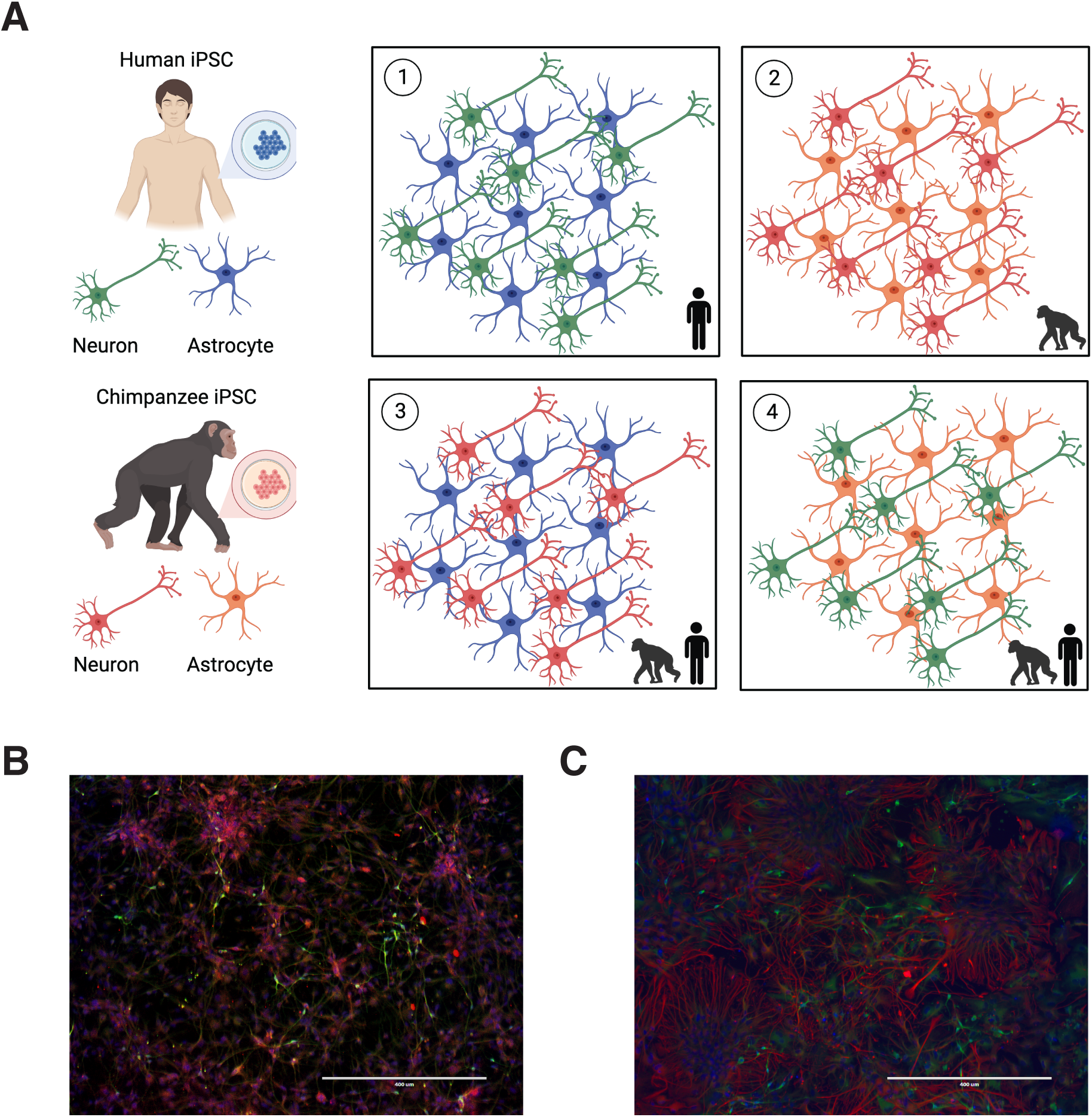
Co-Culture schematic and representative immunofluorescent staining. A: 7 days following differentiation of neurons and astrocytes from both human and chimp-derived iPSCs (see Methods), astrocytes were seeded directly on top of neurons at a 1:1 ratio. Panel A depicts the four co-culture conditions, including two same species (1-2) and two mixed species (3-4). Schematic made in BioRender. B: Immunofluorescent staining of co-cultured human astrocytes and neurons after 6 days (Astrocytes: GFAP (red), Neurons: TUBB3 (green)). 10X magnification. C: Immunofluorescent staining of co-cultured chimpanzee astrocytes and neurons after 6 days (Astrocytes: GFAP (red), Neurons: TUBB3 (green)). 10X magnification.

## Results

### Monoculture metabolic assays recapitulate known cell-type and species differences in metabolism

We first assessed mitochondrial function in living iPSC-derived neurons and astrocytes grown separately in monoculture using the Seahorse Mito Stress Test assay system (Agilent Technologies). Across all parameters surveyed in the Mito Stress Test (basal respiration, maximal respiration, ATP production, and spare respiratory capacity) astrocytes exhibited significantly higher oxygen consumption rates (OCR) than neurons (Figure 2A-C,E-F). This was consistent in human and chimpanzee cells, suggesting that these metabolic differences between neurons and astrocytes are conserved across species.

**Figure 2.**
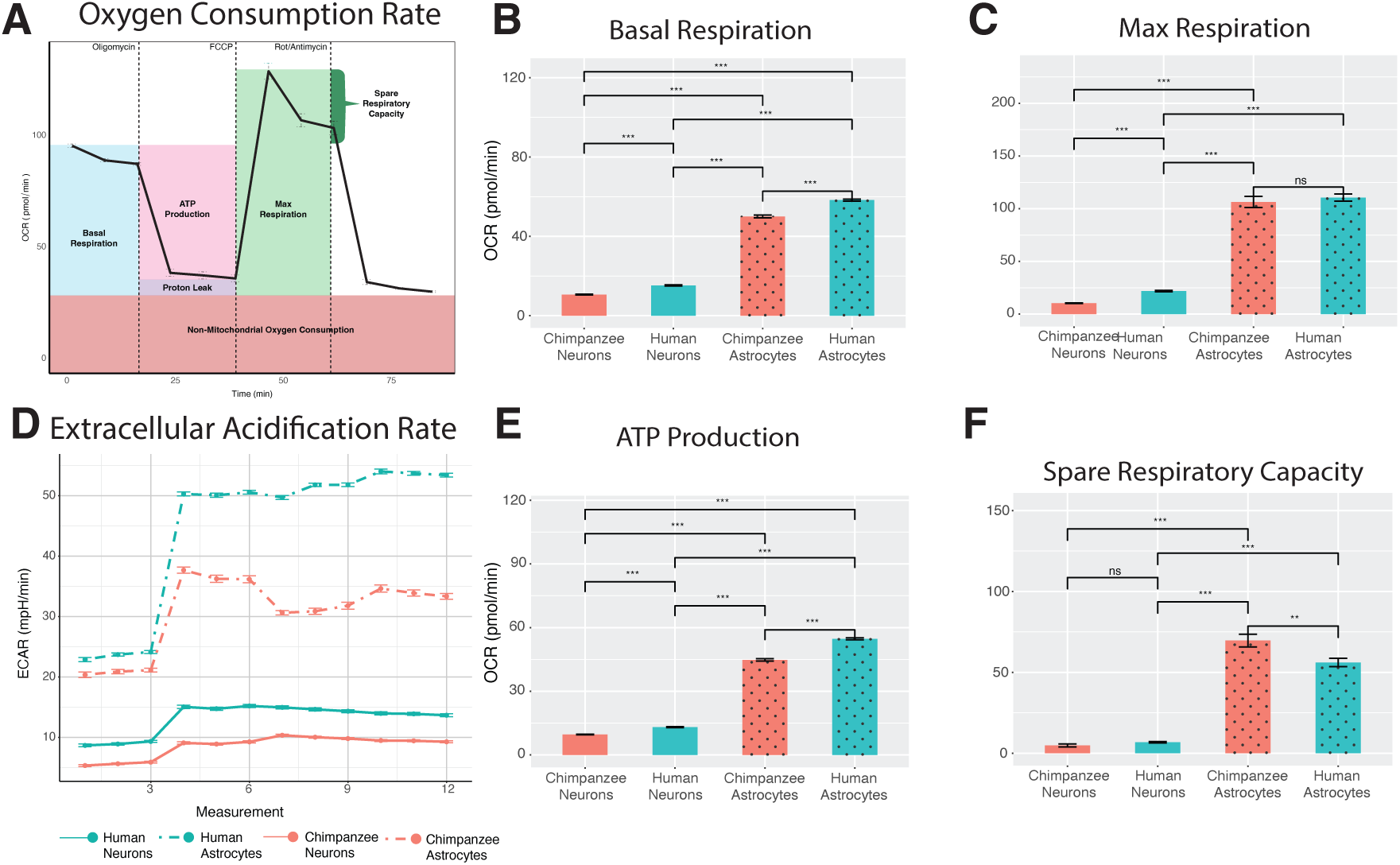
Seahorse MitoStress Test results for monocultured astrocytes and neurons. A: Schematic of the stages of the MitoStress test experiments. B: Boxplots comparing basal respiration rates. C: Boxplots comparing max respiration rates. D: Extracellular acidification rates (ECAR) plotted across the 12 measurements (timepoints) of the experiment). E: Boxplots comparing ATP production. F: Boxplots comparing spare respiratory capacity Error bars represent standard error of the mean (SEM). Significance between groups was assessed using unpaired two-sided t-tests; asterisks denote significance levels (p < 0.05, p < 0.01, p < 0.001, ns = not significant).

Within each cell type, species-specific aspects of metabolic function were observed (Figure 2). Human neurons exhibited significantly increased basal respiration (FDR = 2.26 x 10^-20^), maximal respiration (FDR = 1.04 x 10^-24^), and ATP production (FDR = 8.55 x 10^-22^) (Figure 2 B-C,E-F; Supplementary Table 1) compared to chimpanzee neurons. There was no significant species difference in neuron spare respiratory capacity (FDR = 0.06), signaling that the ability of neurons to respond to energetic stress is evolutionarily conserved.

Human astrocytes similarly exhibited increased basal respiration (FDR = 7.75 x 10^-18^) and ATP production (FDR = 1.87 x 10^-28^) compared to chimpanzee astrocytes (Figure 2B,E-F; Supplementary Table 1). There was no species difference in astrocyte maximal respiration (FDR = 0.51)(Figure 2C; Supplementary Table 1). Interestingly, the chimpanzee astrocytes show a modest increase in spare respiratory capacity compared to the human astrocytes (FDR = 0.0058) (Figure 2F; Supplementary Table 1). This is the only result in which chimpanzee OCR is higher than human OCR, suggesting that chimpanzee astrocytes have a larger metabolic “reserve” that may allow them a more flexible metabolic strategy. In contrast, human astrocytes may be more well-suited to operate closer to their maximal capacity at baseline.

The Mito Stress Test also revealed species- and cell-type-specific differences in extracellular acidification rate (ECAR), a proxy for glycolytic activity. Astrocytes produced significantly elevated ECAR values compared to neurons in both species across all twelve timepoints (all FDR-adjusted q-values < 1.3 × 10^⁻52^) (Figure 2D; Supplementary Table 3). Human astrocytes showed exceptionally high ECAR compared to chimpanzee astrocytes (all FDR-adjusted q-values < 4.72 x 10-5), especially heightened following Oligomycin injection which is consistent with enhanced glycolytic capacity (measurements 3 and 4). Neuron ECAR was much lower overall, a reflection of increased investment in oxidative phosphorylation over glycolysis. Even so, human neurons exhibited significantly higher ECAR than chimpanzee neurons, suggestive of some uniquely human increase in glycolytic output (all FDR-adjusted q-values < 7.13 x 10^-19^). These results ultimately recapitulate known species differences in metabolism, in which the human brain relies more on glycolysis for energy production. In this context, astrocytes exhibit markedly high oxygen consumption rates, suggesting that this cell type plays a key role in shaping the overall metabolic profile of the primate brain, and may contribute to species-specific metabolic strategies.

### Metabolic assays of cells in co-culture highlights species differences in neural cell interactions

We extended our analysis beyond monocultures to examine how interactions between cell types influence overall brain metabolism through a 2D co-culture model of iPSC-derived neurons and astrocytes cultured at a 1:1 ratio. In this experimental setup, we compared same species neural co-cultures as well as mixed species co-cultures to isolate species-driven and cell type-driven differences in metabolism. We refer to the four co-culture combinations as follows: HAHN = human astrocytes with human neurons, CAHN = chimpanzee astrocytes with human neurons, HACN = human astrocytes with chimpanzee neurons, and CACN = chimpanzee astrocytes with chimpanzee neurons (Figure 3).

**Figure 3.**
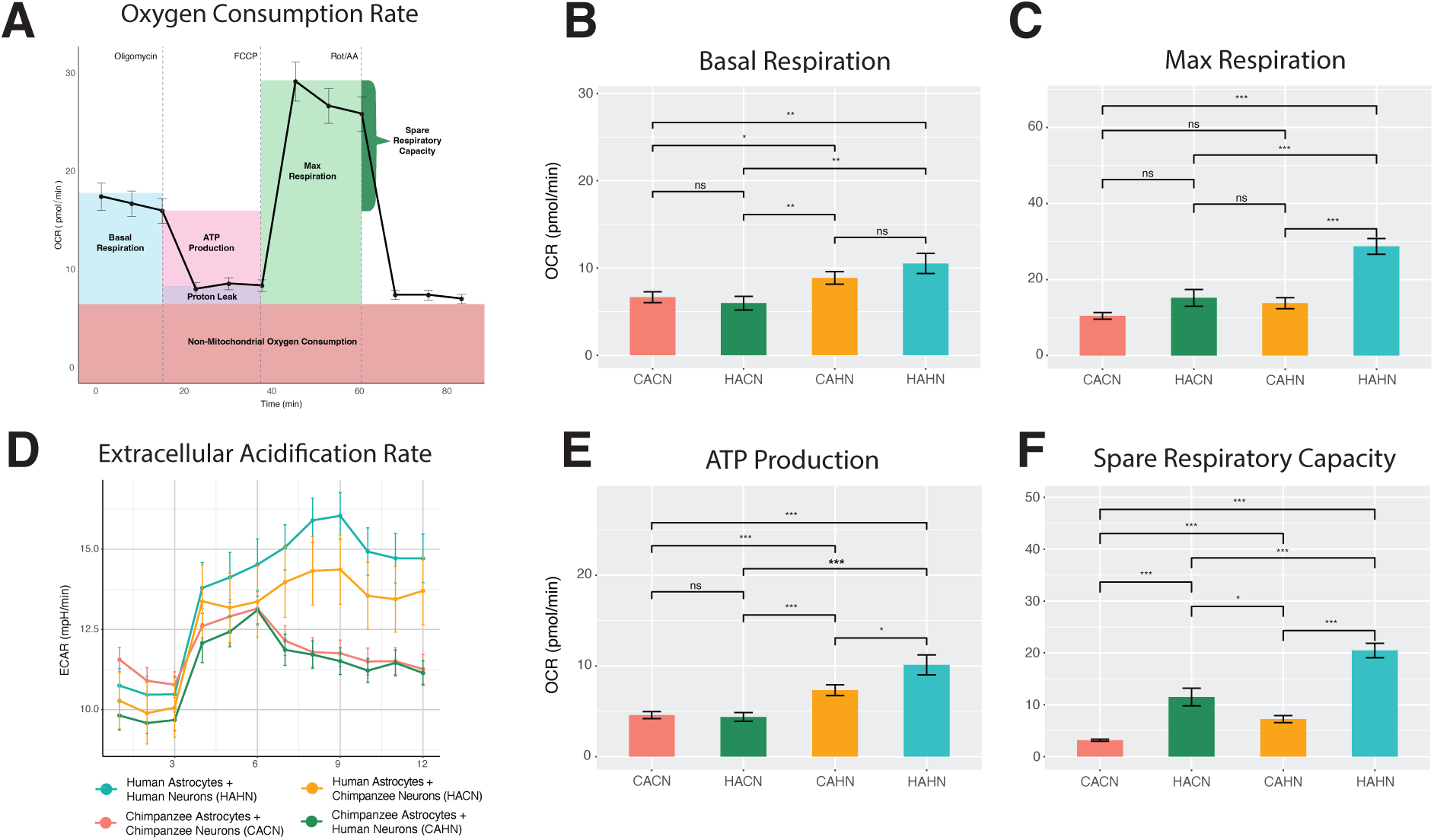
**Seahorse MitoStress Test results for co-cultured astrocytes and neurons**. A: Schematic of the stages of the MitoStress test experiments. B: Boxplots comparing basal respiration rates. C: Boxplots comparing max respiration rates. D: Extracellular acidification rates (ECAR) plotted across the 12 measurements (timepoints) of the experiment). E: Boxplots comparing ATP production. F: Boxplots comparing spare respiratory capacity. Error bars represent standard error of the mean (SEM). Significance between groups was assessed using unpaired two-sided t-tests; asterisks denote significance levels (p < 0.05, p < 0.01, p < 0.001, ns = not significant). Co-culture abbreviations given in 4D.

Similar to the monocultured cells, Seahorse Mito Stress Test data of co-cultured astrocytes and neurons confirmed that human neural co-cultures are more metabolically active than chimpanzee neural co-cultures (Figure 3 B-C,E-F). With this, the human co-culture (HAHN) showed significantly increased basal respiration (FDR = 0.014), maximal respiration (FDR = 6.35 x 10^-9^), spare respiratory capacity (FDR 1.19 x 10^-14^), and ATP production (FDR = 5.79 x 10^-5^) compared to the chimpanzee co-culture (CACN) (Figure 3 B-C,E-F; Supplementary Table 2). We also observed that the human co-culture showed significantly higher ECAR during the later timepoints following treatment (FDR < 0.04), while the chimpanzee co-culture showed higher ECAR during the initial pH measurements preceding treatment (FDR > 0.32) (Figure 3D; Supplementary Table 4). These results indicate greater metabolic capacity in human co-cultures, with stronger glycolytic responses to mitochondrial stress.

The mixed species co-cultures further help to delineate the cell-type specific contributions. Cultures including human neurons (HAHN, CAHN) exhibited higher OCR during basal respiration and ATP production than cultures that include chimpanzee neurons (CACN, HACN) (FDR < 0.03 for all comparisons) (Figure 3B; Figure 3E; Supplementary Table 2). These differences occurred regardless of astrocyte species identity, suggesting that neurons are driving basal respiration and ATP production in the brain.

Cultures including human astrocytes (HAHN and CAHN) exhibited higher OCR during measurements of spare respiratory capacity than cultures that include chimpanzee astrocytes (CACN and CAHN) (FDR < 3.51 x 10^-5^) (Figure 3F; Supplementary Table 2). Similar to above, these differences occurred regardless of neuron species identity and highlight the significant role of astrocytes in responding to metabolic stress within the brain. In particular, human astrocytes appear especially well-equipped to respond to stress.

Interestingly, no single cell type appears to be driving species differences in maximal respiration. In this data, human co-cultured cells overall produce the highest OCR values (HAHN) (FDR < 6.29 x 10^-5^), and no other co-culture combinations (mixed species (HACN and CAHN) as well as chimpanzee only-cultures (CACN)) differ statistically from each other (Figure 3C; Supplementary Table 2). This suggests that species differences in maximal respiration arise though a combination of factors from astrocytes and neurons. Further, this highlights that the interactions between cells may be driving the larger species differences in maximum metabolic capacity.

Finally, we observed that conditions including human astrocytes (HAHN and HACN) exhibited higher ECAR values compared to co-cultures including chimpanzee astrocytes (CACN and CAHN) (Figure 3D; Supplementary Table 4). Notably, HAHN co-cultures demonstrate the highest overall ECAR, significantly exceeding that of HACN co-cultures (Figure 3D; Supplementary Table 4). This pattern underscores the synergistic effect of human astrocytes and neurons on metabolic activity and suggests that these interactions may play a key role in species differences in glycolytic function.

### Integrated scRNA-seq datasets cluster according to cell type identity

We performed single-cell RNA-sequencing for each co-culture condition and integrated these four single-cell datasets into a single, unified dataset (see Methods). Upon integration, we observed a high degree of overlap of cells across co-culture conditions when visualized using Uniform Manifold Approximation and Projection (UMAP) plots, emphasizing that basal processes involved in typical astrocyte and neuron function are conserved among humans and chimpanzees (Figure 4A; Supplementary Figure 1).

**Figure 4.**
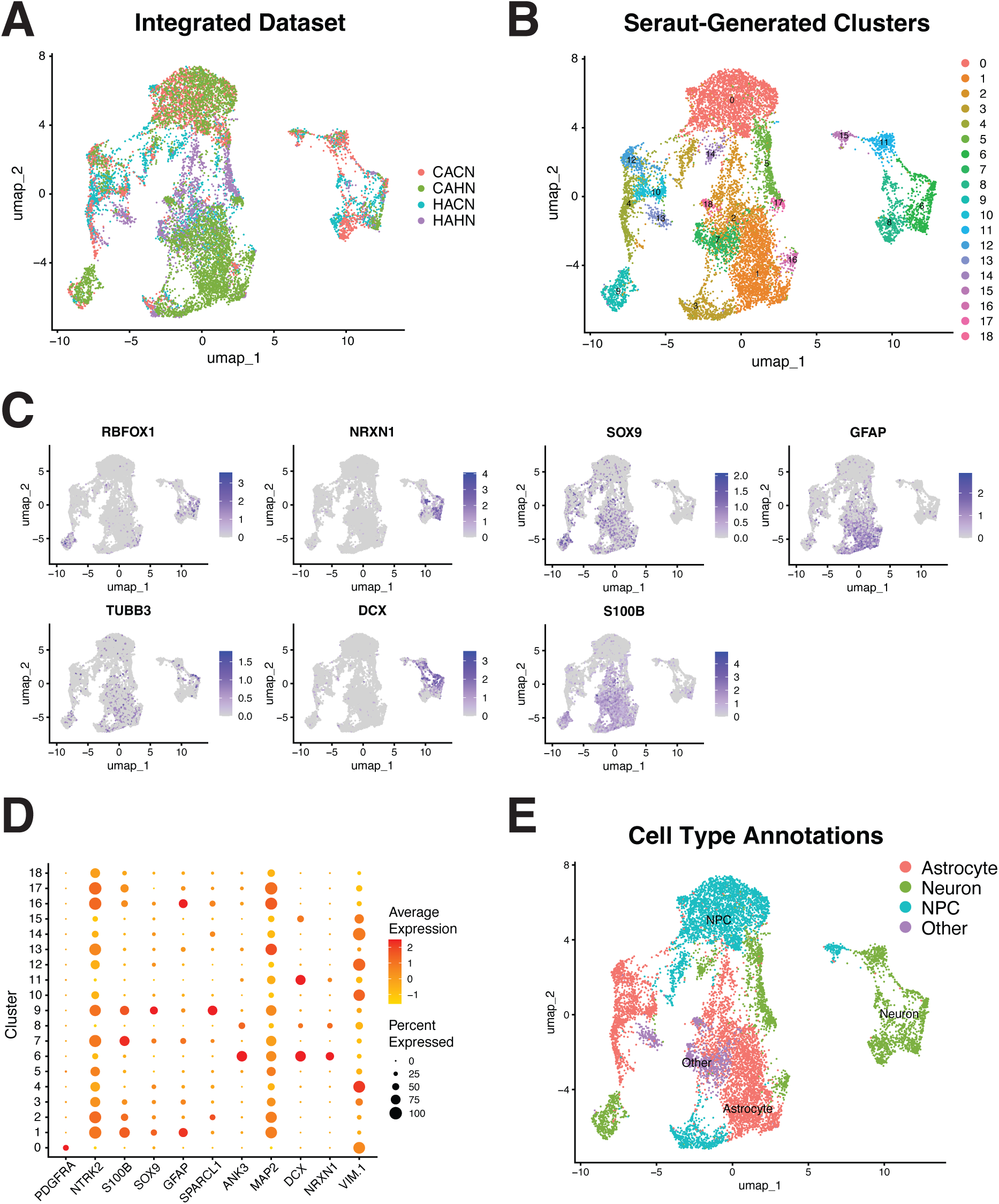
Single cell RNA-sequencing clustering and marker gene expression. A: UMAP of the integrated dataset; cells are colored for which co-culture condition they originated from. B: UMAP of the 19 clusters identified via Seurat. C: Feature plots of neuron (RBFOX1, NRXN1, TUBB3, DCX) and astrocyte markers (SOX9, GFAP, S100B). Cells are colored based on level of expression of marker gene (darker purple = higher expression). D: Dot plots of NPC, neuron, and astrocyte markers by the 19 designated clusters. Color of each dot represents average expression per cluster (more red = higher expression) and the size of the dot represents the percentage of cells in each cluster that express that gene. E: UMAP of integrated dataset annotated by 4 major cell type groupings (Astrocyte, Neuron, NPC, and Other). All plots generated via Seurat in R.

Using the FindClusters function in Seurat (resolution = 0.5), we identified 19 transcriptionally distinct cell clusters (Figure 4B). To assign biological identities to these clusters, we used the FindAllMarkers function in Seraut and focused on expression of established neuronal (i.e. RBFOX1, NRXN1, and DCX) and astrocytic (i.e. GFAP and S100B) marker genes (Figure 4C-D; Supplementary Table 5). Of the 19 communities, 5 were classified as astrocytes (n = 4,653 cells), 8 as neurons (n = 2,887 cells), 3 as immature neural progenitor cells (n = 3,482 cells), and 3 as other minor cell types (792 cells) (Figure 4E). Both the immature neural progenitor cell and other classes were excluded from downstream analysis.

We assigned cluster labels based off marker gene expression and observed that cells generally cluster according to cell-type identity in the integrated UMAP (Figure 4E). This suggests that the major driver of transcriptional variation in this dataset is not species, but rather cell-type identity. Even so, we detected interesting patterns within the neuron and astrocyte clusters that suggest differences across co-culture conditions. For example, Cluster 9 was identified as a neuronal population that is predominantly composed of cells from the CAHN condition (Figure 4A-B; Figure 4E).

This neuron cluster is markedly separated from the other neuron clusters in UMAP space, suggesting that human neuron expression is heavily altered due to the presence of chimpanzee astrocytes. These findings ultimately highlight the intrinsic differences between astrocytes and neurons, while also revealing how gene expression can be altered in the mixed species co-culture conditions.

### Aggregated single-cell data reveals variable expression in genes involved in neural network development and metabolism

We aggregated counts by cell type and co-culture condition (pseudobulk) and normalized for differing cell numbers. We were then able to perform pairwise differential expression analysis using the glm model in EdgeR (see Methods) (Supplementary Table 6; Supplementary Table 8).

We first analyzed differences between the human co-culture (HAHN) and the chimpanzee co-culture (CACN) to establish a baseline for typical interactions that would represent that of a typical human or chimpanzee brain (Supplementary Table 8).

Comparing the human and chimpanzee astrocytes in these same-species co-cultures, we found that the human astrocytes were enriched for genes with functions in “Aerobic Respiration (GO:0009060)”, “Carbohydrate Derivative Metabolic Process (GO:1901135)”, as well as “Oxidative Phosphorylation (GO:0006119)” compared to chimpanzee astrocytes. This suggests that the human astrocytes are genetically primed for a more metabolically active state than the chimpanzee astrocytes when each interacted with neurons of the same species. We also observed that the human neurons had higher expression of genes involved in “Regulation of Primary Metabolic Process (GO:0080090)”, “Regulation of Signaling (GO:0023051)”, “Microtubule-based Process (GO:0007017)” as well as “Glucose Metabolic Process (GO:0006006)” compared to the chimpanzee neurons. This emphasizes that the human neurons are not only more metabolically active than the chimpanzee neurons but are also more primed for communicative signaling between cells. This uniquely human enrichment for signaling-related genes further implies that the human co-cultures may be capable of producing more complex neural networks, as increased signaling capacity supports greater synaptic connectivity and network integration ^34^.

To elucidate which cell type is driving species differences in our co-culture model, we next investigated pairwise differences in gene expression between same and mixed species co-culture conditions (Supplementary Table 8). Human astrocytes upregulated genes involved in “Cell Communication (GO:0007154)”, “Postsynaptic actin cytoskeleton organization (GO:0098974)”, and “Neuropeptide signaling pathway (GO:0007218)” in the context of human neurons. Conversely, human astrocytes upregulated genes involved in “Generation of neurons (GO:0048699)”, “Nervous system development (GO:0007399)”, and “Neuron differentiation (GO:0030182)” in the context of chimpanzee neurons.

As with the astrocytes, we also found significant differences in neuronal gene expression between same-species and mixed-species co-cultures (Supplementary Table 8). Human neurons cultured with human astrocytes upregulated genes involved in “Regulation of membrane potential (GO:0042391)”, “Neurotransmitter transport (GO:0006836)” and “Secondary metabolic process (GO:0019748)”. In contrast, when human neurons interacted with chimpanzee astrocytes, upregulated genes instead included “Extracellular matrix organization (GO:0030198)” and “Cell migration (GO:0016477)”. This signals that the human neurons exhibit signatures of increased metabolic and electrochemical activity only when supported by human astrocytes.

Chimpanzee neurons also appear to respond positively to the presence of human astrocytes: chimpanzee neurons in mixed-species cultures were enriched for “Positive regulation of primary metabolic process (GO:0009893)” as well as “Plasma membrane bounded cell projection assembly (GO:0120031)”. This indicates that human astrocytes may alter the culture environment in ways that cause the chimpanzee neurons to increase their metabolic activity, potentially priming them for increased synaptic signaling and more complex neural network formation.

### Global analysis of metabolic gene expression highlights astrocytes as drivers of metabolic differences across humans and chimpanzees

We also examined expression of select metabolism-related gene sets across grouped cells from each co-culture condition. To identify global differences, we performed an ANOVA-like likelihood ratio test using a generalized linear model with a fixed dispersion of 0.2 (Supplementary Table 7). This approach allowed us to detect genes that were differentially expressed across any of the experimental groups.

Notably, key glycolysis enzymes including Hexokinase 2 (HK2), Phosphofructokinase (PFKM; PFKL), Aldolase (ALDOA; ALDOB; ALDOC), and Enolase (ENO2; ENO3) were significantly highly expressed in the cells from the HAHN and HACN conditions, suggesting elevated glycolytic activity when human astrocytes were present (Figure 5; Supplementary Table 7). Importantly, for several of these genes there was no significant difference in expression between the human astrocytes from the HAHN and HACN conditions, suggesting that these changes are not dependent on the species identity of the neurons. Rather, astrocytes appear to be driving high glycolytic rates both within the astrocytes and in the neurons that they interact with.

**Figure 5.**
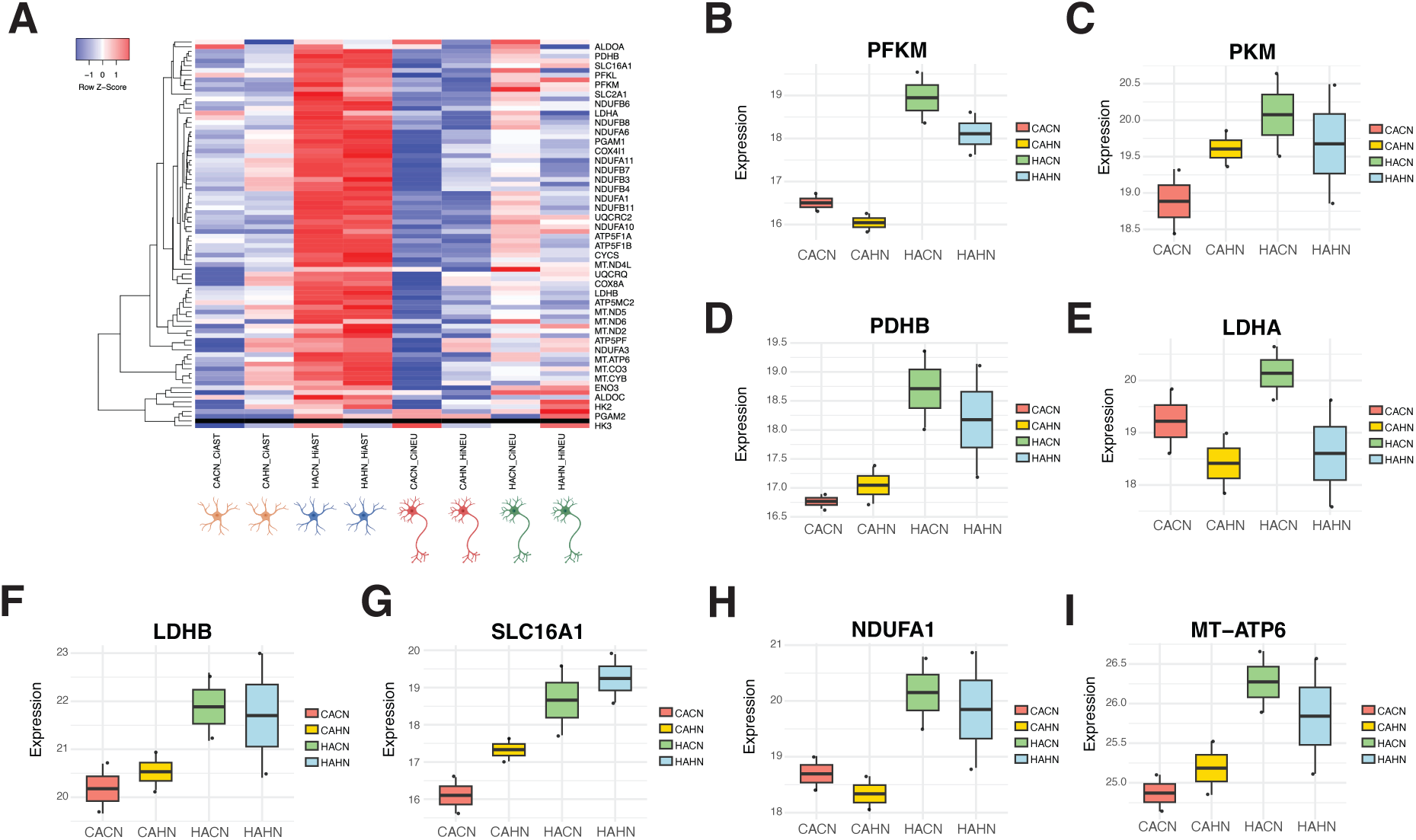
Global differential expression analysis of metabolism-related genes. A: Heatmap of expression of metabolism-related genes separated by grouping. Each row represents a gene and each column a different group of cells from the four co-culture conditions (HAHN = human astrocytes with human neurons; CACN = chimpanzee astrocytes with chimpanzee neurons; HACN = human astrocytes with chimpanzee neurons; CAHN = chimpanzee astrocytes with human neurons; CiAst = chimpanzee astrocytes; HiAst = human astrocytes; CiNeu = chimpanzee neurons; HiNeu = human neurons). Illustrations of cell type are colored by species (orange = chimpanzee astrocytes; blue = human astrocytes; red = chimpanzee neurons; green = human neurons). Heatmap colors represent relative expression levels (red: more highly expressed; blue: more lowly expressed). All heatmaps were generated using heatmapper.ca B-I: Boxplots of specific metabolism-associated genes. Each box represents a co-culture condition (including data from both cell types). The boxplot displays the interquartile range (IQR), with the median indicated by the central line and whiskers extending to 1.5× IQR. Outliers were omitted from the boxplot visualization but are included in the jittered points.

Similarly, genes involved in lactate production and transport (LDHA, PDHB, SLC16A1/4) were elevated in human astrocytes from the HAHN and HACN conditions, pointing to increased aerobic glycolytic flux and mitochondrial pyruvate processing that is again not dependent on neuron identity (Figure 5; Supplementary Table 7) ^35^ ^36^. LDHB, which converts lactate into pyruvate for oxidative metabolism, also showed increased expression in the human astrocytes from these conditions, however these results were not statistically significant (Figure 5A; Figure 5F; Supplementary Table 7). Ultimately, these results highlight that the human astrocytes have likely evolved a more efficient and potentially more active lactate shuttling mechanism.

Finally, we observed widespread upregulation of electron transport chain (ETC) genes in conditions containing human astrocytes, suggesting enhanced mitochondrial respiration (Figure 5A; Figure 5H; Supplementary Table 7). For example, UQCRC1, which is involved in the formation of Complex III, was expressed significantly higher in the human-astrocyte containing groups relative to conditions containing chimpanzee astrocytes ^37^. Other genes that encode subunits of ATP Synthase (MT-ATP6; MT-ATP8) as well as those that encode Cytochrome c Oxidase (MT-CO1; MT-CO2) were also expressed significantly higher in conditions containing human astrocytes (Figure 5A; Figure 5I; Supplementary Table 7). This pattern suggests that human astrocytes may promote increased mitochondrial transcription both intrinsically and in neighboring neurons, contributing to enhanced oxidative function of the neural cell network.

## Discussion

In this study, we investigated species level differences in the metabolic activity of neurons and astrocytes, and further how interactions between these cell types alter metabolic states in a co-culture model.

First, in our analysis of monocultured iPSC-derived neurons and astrocytes, we were able to distinguish cell-type and species-specific changes in oxygen consumption using the Seahorse XF Mito Stress Test. Compared to neurons, astrocytes exhibited higher overall metabolic output – highlighting their significant functional role as the major energetic support system for the brain. This reinforces previous transcriptomic data collected from human and chimpanzee iPSC-derived neural progenitor cells, neurons, and astrocytes ^26^. We acknowledge that the iPSC-derived neurons and astrocytes are differentiated in cell-type specific medias and thus may be subject to differing glucose concentrations prior to the Seahorse Assay. However, both cell types are similarly treated with Seahorse XF DMEM medium one hour prior to assay, which should help to normalize any acute effects from the culture medium.

We also observed that human neural cells (both neurons and astrocytes) had higher basal respiration rates and higher ATP production than the chimpanzee neural cells. Only in the neurons did we detect a significant species difference in max respiration rates and spare respiratory capacity, suggesting that the human and chimpanzee astrocytes may be functioning more similarly than the neurons in monoculture.

Seahorse assays from co-cultured cells further revealed how interactions between these neurons and astrocytes impact the overall metabolic output of the neural network. We confirmed that human neural cells were more metabolically active than chimpanzee cells, both in the monoculture and in the co-culture, supporting data from living brain tissue in which humans were found to consume roughly 20% of the body’s resting metabolic resources compared to only 10% in chimpanzees ^1–3,14,38^.

Our cross-species co-cultures allowed us to separate out the cell-type specific effects in these interaction models. We conclude that neurons are likely drivers of basal metabolic rates as well as ATP production. This is likely a reflection of neurons favoring oxidative phosphorylation as their dominant metabolic process ^26,38,39^. Importantly, this also provides a potential mechanism for the evolution of the unique complexity of the human brain: neurons have evolved to produce ATP at a more rapid rate in the human brain, resulting in the energy necessary to facilitate synaptic connections that ultimately lead to complex neural network formation ^40,41^.

In contrast to neurons, astrocytes in the co-culture model drive differences in spare respiratory capacity. This data reflects known functional differences between astrocytes and neurons in which astrocytes can store large amounts of glycogen within themselves, whereas neurons cannot ^42–44^. With this, it becomes more efficient for the brain to house energy stores within the astrocytes and shuttle those nutrients to the neurons when needed ^43,45^. This also highlights astrocytes as key regulators of homeostasis within the brain, and more specifically how these cells can upregulate oxidative metabolism in response to stressful or energetically demanding conditions ^46^. In particular, human astrocytes are especially well-equipped with energetic reserves, allowing them greater flexibility than chimpanzee astrocytes in our co-culture system.

Extracellular acidification rates (ECAR) across the mixed and same-species co-culture conditions underscore how humans and non-human primates may differ in the astrocyte-neuron lactate shuttle (ANLS). ECAR is a measure of the change in pH of the culture environment and an increase in ECAR has been noted to be caused by increases in lactate production following glycolysis ^47^. Taking this into consideration, our data illustrates that human co-cultures produce higher ECAR than chimpanzee co-cultures, and the cross-species conditions further reveal that the human astrocytes are likely driving these increases in ECAR. This ultimately suggests that human astrocytes produce high amounts of lactate, which can then be shuttled to the neurons via the ANLS ^18,22^. Importantly, lactate conversion into pyruvate does not require glycolysis (a slow, enzyme-intensive process), and so neurons may preferentially utilize this lactate delivered from the astrocytes as a faster means to ATP synthesis ^48^. From there, the neurons can process this excess lactate into ATP via oxidative metabolism, which can then be used to power synaptic connections and facilitate neural network formation ^49^.

Importantly, lactate production via astrocytes may serve additional functions outside of energy production that further explain some of the species-differences in primate brain evolution. For example, increased lactate levels have been found to enhance calcium influx via NMDA receptors and influence downstream synaptic plasticity ^50^. In addition, lactate has been implicated in modulating immune responses and promoting resistance to oxidative stress ^50,51^. Thus, lactate may not only support neuronal energy demands but also directly regulate neural signaling and resilience - processes that may influence susceptibility to disease ^49^.

Single cell sequencing data from these multi-species co-cultures further corroborates much of the data from the live cell Seahorse Metabolic Assays. Differential gene expression analysis confirmed that human astrocytes show signatures of enhanced glycolytic capacity and metabolic flexibility relative to the chimpanzee astrocytes. With this, human astrocytes were uniquely enriched in genes involved in enhanced glycolysis, lactate shuttling, as well as oxidative phosphorylation. This enhanced metabolic activity was consistent regardless of the species identity of the neurons, suggesting that human astrocytes are highly adaptive to the surrounding environment and are similarly able to support human and chimpanzee neurons.

Expression data also emphasized how neurons are metabolically responsive to astrocyte identity. For example, in the context of human astrocytes, the chimpanzee neurons upregulated the expression of several oxidative metabolism related genes, suggesting that they are capable of higher metabolic output but are likely limited by the chimpanzee astrocytes. Human neurons were also found to strongly respond to the chimpanzee astrocytes, downregulating oxidative metabolism and lactate shuttling genes. This further emphasizes that chimpanzee astrocytes provide limited metabolic support, which forces human neurons to dampen their metabolic activity. Together, these results suggest that species-specific differences in astrocyte function can directly influence neuronal energy metabolism, potentially contributing to the distinct metabolic profiles observed across the whole primate brain.

Single-cell sequencing data also revealed interesting patterns of cell-type densities within the different co-culture setups. We observed a substantial number of cells denoted as neural progenitor cells (NPCs) in the chimpanzee co-culture (CACN) and the mixed chimpanzee astrocyte-human neuron co-culture (CAHN) (Figure 5A-B). This suggests that the chimpanzee astrocytes may represent a more immature stage of astrocyte differentiation relative to the human astrocytes. We also were able to distinguish populations of specific neuron subtypes – for example some expressing excitatory (Cluster 8) marker genes like NRXN1, ANK3, GRIK2, and ANKS1B (Supplementary Table 5). Future analysis of the relative ratios of excitatory to inhibitory subtypes in each co-culture may aid in our understanding of the supportive roles of astrocytes and how these might differ across primates.

## Conclusion

Our novel cross-species co-culture approach reveals hidden metabolic plasticity and intercellular communication differences that are indiscernible within a single-species monoculture or co-culture system. Whereas 2D monocultures provide information on cell-intrinsic glycolytic capacities, co-cultures better characterize the interactions that exist within living brain tissue. These co-cultures are also capable of enhancing the maturation of both the astrocytes and neurons, which also contributes to a better model of post-embryonic brain tissue. In summary, astrocytes are highly flexible neural cells that drive major species differences in brain metabolism. Neurons, while responsible for forming the synaptic architecture of the brain, rely heavily on astrocyte support to regulate their activity and guide the maturation of neural networks. This data underscores the critical role of astrocyte–neuron interactions in shaping brain evolution and further highlights that pathways underlying these interactions (such as the ANLS) may be key contributors to both human-specific brain function and disease vulnerability.

## Methods

### Cell lines and maintenance

We utilized two human (Hi28834 and Hi20682, both female) and two chimpanzee (Ci3647B1 and Ci40210, both female) iPSC cell lines for this cross-species study.

These iPSCs were previously generated from fibroblasts collected from minimally invasive skin biopsies, and following reprogramming all iPSCs were validated for pluripotency and lack of any karyotype abnormalities ^52–58^. These feeder-free iPSCs were maintained in mTeSR1 media (STEMCELL Technologies, Vancouver, Canada) on Matrigel-coated plates (Corning, Keene, NH, USA). All iPSCs were cultured for at least 2 passages prior to differentiation.

### Differentiation into astrocytes and neurons

We used established protocols from STEMCELL technologies to differentiate iPS cells into neurons and astrocytes. We first induced cell lines into Neural Progenitor Cells (NPCs) using Neural Induction Medium + SMADi supplement for a minimum period of 18 days on Matrigel-coated plates (STEMCELL Technologies; Corning). To differentiate astrocytes, we cultured NPCs in Astrocyte Differentiation Media on Matrigel-coated plates for a period of 21-28 days, followed by culture in Astrocyte Maturation Media for a period of at least 21 days (a minimum of 6 weeks to mature astrocytes). To differentiate forebrain neurons, we cultured NPCs in Forebrain Neuron Differentiation Media for a period of 6-8 days, followed by culture in BrainPhys Media with Forebrain Neuron Maturation Supplements for 7-14 days (a minimum of 2 weeks to mature neurons).

Neurons were grown on Poly-L-Ornithine(PLO)/Laminin-521 (Millipore Sigma #P4957; #L2020) coated plates following protocols from StemCell Technologies.

All differentiations were verified through immunofluorescent staining of cell-type specific markers. For each culture, iPSCs were fixed in 4% Paraformaldehyde in PBS for 10 minutes. NPCs and Astrocytes were permeabilized with 0.3% Triton X-100 for 7 minutes. Neurons were similarly permeabilized with 0.1% Triton X-100 for 7 minutes. Samples were then blocked for 60 minutes with 3% Bovine Serum Albumin (BSA) in PBS at room temperature (RT). Primary antibody solutions were diluted in 3% BSA and incubated overnight at 4°C. Secondary antibody solutions were similarly diluted in 3% BSA and incubated for 45 minutes at RT. Finally, samples were mounted with ProLong Gold Antifade Mountant with DAPI (Thermo Fisher Scientific). The following cell type-specific markers were utilized for the respective cell types: NPCs for PAX6+/OCT4-, neurons for β-Tubulin III+/PAX6-, and astrocytes for GFAP+/PAX6-.

### Co-Culture of iPSC-derived astrocytes and neurons

Co-cultures were set up following successful neuron differentiation on PLO-Laminin coated plates using the Hi28834 and Ci3647B1 cell lines. First, iPSC-derived GFAP+ astrocytes were dissociated from monoculture and resuspended in Astrocyte Maturation Medium. Astrocytes were seeded directly onto the neuron monoculture (without passaging neurons) at a 1:1 ratio. Other tested ratios included 1.5:1 and 2:1 with no significant differences in cell death. A 1:1 ratio was ultimately selected for downstream assays, consistent with recent estimates of glia:neuron ratios in the human brain and comparable regions in non-human primates ^13,59,60^. Wells designated for single-cell sequencing were seeded at a density of 600,000 neurons and 600,000 astrocytes, whereas wells designated for immunofluorescent staining were seeded at a density of 300,000 neurons and 300,000 astrocytes (in a 6-well plate). Seeding densities used in the Seahorse Assay are described below. After 24 hours, the Astrocyte Maturation media was replaced with Neuron Maturation Medium (BrainPhys + neuron maturation supplement, STEMCELL Technologies). Co-cultures were maintained for 48-hours prior to Seahorse Assays and 8 days prior to single cell isolation (with media changes every 2 days).

We included single-species and mixed species co-cultures for 4 total conditions: (1) human astrocytes with human neurons, (2) chimpanzee astrocytes with chimpanzee neurons, (3) human astrocytes with chimpanzee neurons, and (4) chimpanzee astrocytes with human neurons (Figure 1).

### Seahorse Assay prep and plate seeding

We utilized the Seahorse Cell Mito Stress Test Kit to measure changes in metabolic rate across cell types and co-culture conditions (Figure 3-4A) (Agilent Technologies, Santa Clara, CA). Assays were performed on both monocultured and co-cultured astrocytes and neurons. For the monocultures, 15,000 cells were seeded into wells of a Seahorse XF96 plate (Agilent Technologies, Santa Clara, CA). For the co-cultures, 7,500 cells each of astrocytes and neurons were seeded (total = 15,000 cells) in XF96 plates.

In the Seahorse experiments involving neurons (monoculture and co-culture), neuron precursor cells were seeded into XF96 plates 6–7 days after initiating neuronal differentiation and maintained for an additional 10 days in Neuron Maturation Media to promote maturation and expression of TUBB3. Monocultured astrocytes were seeded into XF96 plates after reaching mature GFAP expression, and assays were conducted 48 hours after initial seeding. For co-cultures, following neuron maturation and confirmation of TUBB3 expression, GFAP+ astrocytes were seeded directly into wells containing neurons of one of the two species (see protocol above).

Human and chimpanzee cells were seeded in the same plates to limit batch effects. However, monocultured neurons and astrocytes were seeded in separate plates due to differences in differentiation protocols (neurons have to mature in the Seahorse plate whereas astrocytes can be passaged following maturation) as well as coating differences (PLO-Laminin for neurons, Matrigel for astrocytes). Equal counts of wells were used for the human and chimpanzee conditions. Four wells with no cells were set as background control in all analyses.

Assay medium was prepared prior to the assay, consisting of 10 mM glucose, 1 mM pyruvate, 2 mM glutamine, and Seahorse XF DMEM medium, pH 7.4. (Agilent Technologies, Santa Clara, CA). XF calibrant was also added to the sensor cartridge following overnight incubation of the cartridge with sterile water in a 37 ℃ non-CO2 incubator.

### Seahorse Cell Mito Stress Test Assay

The Seahorse XF sensor cartridge, pre-filled with XF calibrant solution, was incubated for one hour at 37 °C in a non-CO₂ incubator to allow for equilibration. Cells were washed with pre-warmed assay medium, and cells were incubated in 180 µL of assay medium per well for one hour in a 37 °C non-CO₂ incubator to permit equilibration to Seahorse assay conditions. Following incubation of the sensor cartridge, the following drug treatments were loaded sequentially into the appropriate ports of each well: 20 µL of Oligomycin (1.5uM), 22 µL of FCCP (2.0uM), and 25 µL of Rotenone/Antimycin A (0.5uM each). Each of these drug treatments targets a different aspect of the Electron Transport Chain (ETC). Oligomycin inhibits ATP synthase, allowing for indirect measurements of ATP-linked respiration. FCCP acts as an uncoupling agent in the ETC, and its addition reveals the maximal respiratory capacity of the culture system. Finally, the combination treatment of Rotenone and Antimycin A inhibits complexes I and III of the ETC and fully inhibits all mitochondrial respiration, allowing us to estimate non-mitochondrial respiration rates.

Finally, the loaded cartridge was inserted into the Seahorse XFe96 Analyzer (Agilent Technologies, Santa Clara, CA) for calibration. Cells were washed again and 180uL of fresh assay medium was added into each well before loading the assay plate into the Seahorse Analyzer. The Seahorse Analyzer measures Oxygen Consumption Rate (OCR), Extracellular Acidification Rate (ECAR), and Proton Efflux Rate (PER) in each well every 6.5 minutes for a total of 12 measurements (about 80 minutes).

Oligomycin is injected into the assay plate wells after the third measurement, FCCP after the sixth measurement, and Rotenone/Antimycin A after the ninth measurement.

### Seahorse Assay data processing and analysis

Seahorse XFe96 Cell Mito Stress Test assay data were exported from Wave software (Agilent Technologies) as Excel files and analyzed in R (v4.2.0). Raw oxygen consumption rate (OCR) values were read using the readxl package, and missing values were removed using na.omit(). Experimental conditions were identified by the Group variable, and data from individual wells were subset accordingly to reflect the four monoculture conditions and the four co-culture conditions (8 groups total).

Monoculture and co-culture data were compared separately.

Raw OCR values were separated by timepoint (or “Measurement,” corresponding to 1–12) for downstream calculations in each well. Non-mitochondrial oxygen consumption was calculated as the minimum OCR after Rotenone/Antimycin A treatment (measurements 10-12). To estimate basal respiration, the non-mitochondrial respiration rate was subtracted from the last OCR measurement pre-injection (measurement 3). Max respiration was determined by subtracting the non-mitochondrial respiration rate from the maximum OCR after FCCP injection (measurements 7-9). ATP production was estimated as the difference between the basal OCR (measurement 3) and the minimum OCR following oligomycin injection (measurements 4–6). Spare respiratory capacity, an indirect measurement, was calculated as the difference between maximum respiration and basal respiration. To improve data quality, outliers were iteratively removed from each metric using a custom IQR-based filtering function, identifying data points 1.5X outside the interquartile range. Only wells with valid, positive OCR values were retained.

Mean and standard error values for basal respiration, maximum respiration, spare respiratory capacity, and ATP production in each group were visualized as bar plots using ggplot2 (Figure 2-3; Supplementary Table 9). To assess statistical significance between groups, pairwise t-tests were performed using the t.test() function (Supplementary Tables 1-2). For Extracellular Acidification Rate (ECAR), we similarly averaged and calculated standard error values for each group over all 12 timepoint measurements (Figure 2-3; Supplementary Tables 3-4; 9).

### Single Cell Sequencing

Astrocytes and neurons were maintained in co-culture for 7 days prior to obtaining single cell suspensions. Cells were harvested via dissociation with 0.25% Trypsin-EDTA (Gibco #25200056, Thermo Fisher Scientific, Waltham, MA) and filtered through a 40 um cell strainer to remove cell aggregates (Millipore Sigma #CLS431750-50EA, Burlington, MA). Following cell harvesting, cells were washed with 0.04% BSA in 1X PBS (Thermo Fisher #AM2616; #10010023, Waltham, MA) (following the 10X Genomics Sample Prep Demonstrated Protocol for Cultured Cells). Prior to cell counting, the cell suspensions were filtered again through a 40 um cell strainer. Cell counts and viability were estimated with AO/PI staining via the Nexcelom Bioscience Cellometer K2 (Nexcelom Bioscience, Lawerence, MA). Samples were all at least 80% viable and we targeted 5,000 cells/sample for sequencing.

Single cell suspensions were loaded onto the 10X Genomics Chromium Controller using the Chromium Next GEM Chip G and corresponding reagents (Gel Beads and Partitioning Oil) following 10X Genomics guidelines (10X Genomics, Pleasanton, CA). Individual cells, barcoded gel beads, and master mix were co-encapsulated in Gel Bead-In-Emulsions (GEMs). Following GEM generation, an RT incubation step allowed for generation of barcoded cDNA within the GEMs. Following this, emulsions were broken, and cDNA was purified using Dynabeads MyOne SILANE beads. cDNA quality and quantification was assessed using the Agilent 21000 Bioanalyzer High Sensitivity system (Agilent Technologies, Santa Clara, CA).

Libraries were prepared according to manufacturer’s instructions and uniquely indexed (10X Genomics Chromium Next GEM Single Cell 3’ Kit v3.1 User Guide #CG000204 Rev. D). Libraries were quantified via the Agilent 2100 Bioanalyzer High Sensitivity system prior to sequencing (Agilent Technologies, Santa Clara, CA).

Barcoded libraries were sequenced by the Novogene Corporation staff on the llumina NovaSeq X Plus at a depth of at least 50Gb, corresponding to 100 million paired end reads and roughly 20,000 reads per cell.

We created a custom reference sequence that combines human and chimpanzee reference genome sequence and annotations using the mkref function in CellRanger ^61^. For this combined reference, we concatenated Ensembl reference genomes and GTF annotation files from both species (GRCh38 for humans, Pan_tro_3.0 for chimpanzee). Raw reads were first processed using the CellRanger count function with default settings, which aligns reads to the reference and generates a gene-by-cell count matrix. Matrices for each sample were imported into R (v4.3.1) and processed using the Seurat package (v5.0.1) ^62–66^. We assigned metadata to samples based on astrocyte species, neuron species, culture type (mixed or same species), and co-culture condition (HAHN, CACN, HACN, CAHN). Cells were filtered to remove low-quality barcodes based on mitochondrial gene content (>10%), gene detection (<100 or >8,000 features), and UMI counts (<500 or >35,000). We normalized samples using the SCTransformation function, which normalizes, stabilizes variance, and removes technical effects ^62–66^.

Datasets were integrated using Canonical Correlation Analysis (CCA) via the IntegrateLayers() function, and the RNA assay layers were joined. Clustering was performed using the first 30 dimensions, and a resolution of 0.5 yielded 19 distinct communities. Marker genes for each cluster were identified using FindAllMarkers(), and visualization was performed using DimPlot(), FeaturePlot(), and DotPlot() (Supplementary Table 5).

Cell type identities were assigned to clusters based on the expression of canonical marker genes for neurons (e.g., RBFOX3, MAP2, SNAP25) and astrocytes (e.g., GFAP, S100B, ALDH1L1). For downstream analysis, pseudobulk gene expression matrices were generated by aggregating raw UMI counts across cells within each cluster, grouped by both inferred cell type and species of origin as annotated in sample metadata. Differential expression analysis was performed on these pseudobulked datasets using the edgeR package (v3.42.4) ^67^. Genes with low counts were filtered, and library sizes were normalized using the trimmed mean of M-values (TMM) method. We also performed a log-transformed offset from the cell counts for each group and applied this to the model in order to account for differences in the number of contributing cells per pseudobulked sample. Log-transformed counts per million (logCPM) were computed from the normalized data and a design matrix was constructed based on co-culture condition, cell type, and species identity (resulting in 8 total groups) (Supplementary Table 6). A negative binomial generalized linear model was fit to the data using glmFit(), and statistical testing was conducted using the likelihood ratio test via the glmLRT() function (Supplementary Tables 7-8). We assumed a constant biological dispersion of 0.2, reflecting relatively moderate variation in our dataset. An FDR-corrected p-value threshold of <0.05 was utilized to rank significant differences in expression. However, because pseudobulk samples were aggregated without true biological replicates per condition, these FDR-corrected p-values should be interpreted with caution. Unsupervised biological process enrichment analysis was performed using differentially expressed genes (DEGs) using g:Profiler’s functional enrichment tool (g:GOSt) ^68^. Enrichments with a q-value of <0.05 were considered as significant.

## Acknowledgements

We would like to acknowledge our funding from NSF BCS-1750377 and a fellowship to KR from NIH T32 GM135096. We would also like to thank the Genomics Core Facility at UMass Amherst.

## Data availability

Sequencing data have been deposited in the Short Read Archive as: https://dataview.ncbi.nlm.nih.gov/object/PRJNA1443372?reviewer=a7g51aad48m8o097iii1d94b2r And https://dataview.ncbi.nlm.nih.gov/object/PRJNA1440420?reviewer=the4c1hn5l8ob926c4mr6se6lu

---

**Supplementary Figure 1:**
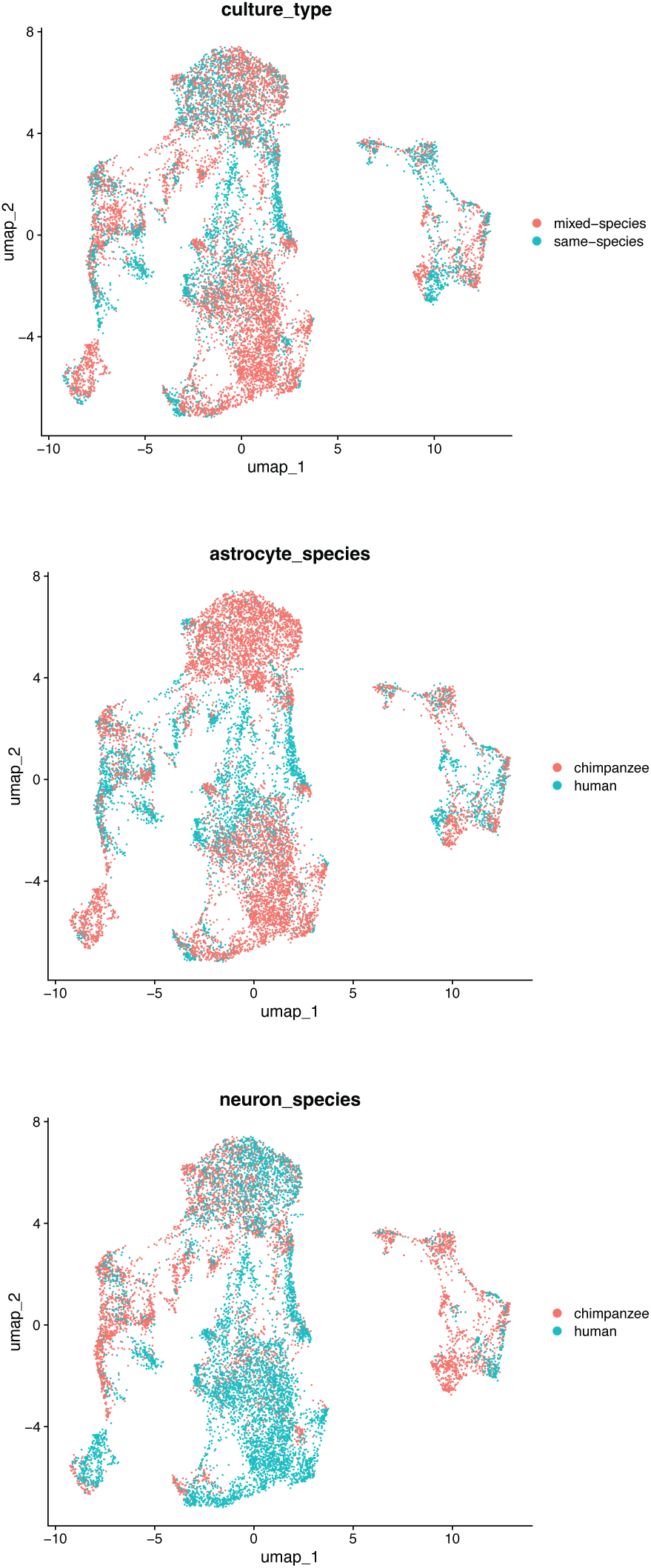
UMAP of integrated single cell RNA-seq datasets, colored based on culture type (A) astrocyte species (B) or neuron species (C) (from assigned sample metadata). UMAPs do not include cell-type specific annotations. UMAPs were generated via Seurat in R using the integrated scRNA-seq dataset (see methods).

## Notes

### Competing Interest Statement

The authors have declared no competing interest.

